# Translation regulation of Japanese encephalitis virus revealed by ribosome profiling

**DOI:** 10.1101/2020.07.16.206920

**Authors:** Vaseef A. Rizvi, Maharnob Sarkar, Rahul Roy

**Affiliations:** Molecular Biophysics Unit; Department of Chemical Engineering, Indian Institute of Science, Bengaluru, India

**Keywords:** Japanese encephalitis virus, translation, ribosomal frameshifting, tRNA, upstream ORF, neuroinvasiveness

## Abstract

Japanese encephalitis virus (JEV), a neurotropic flavivirus, is the leading cause of viral encephalitis in endemic regions of Asia. Although the mechanisms modulating JEV virulence and neuroinvasiveness are poorly understood, several acquired mutations in the live attenuated vaccine strain (SA14-14-2) point towards translation regulation as a key strategy. Using ribosome profiling, we identify multiple mechanisms including frameshifting, tRNA dysregulation and alternate translation initiation sites that regulate viral protein synthesis. A significant fraction (~ 40%) of ribosomes undergo frameshifting on NS1 coding sequence leading to early termination, translation of NS1′ protein and modulation of viral protein stoichiometry. Separately, a tRNA subset (glutamate, serine, leucine and histidine) was found to be associated in high levels with the ribosomes upon JEV infection. We also report a previously uncharacterised translational initiation event from an upstream UUG initiation codon in JEV 5′ UTR. A silent mutation at this start site in the vaccine strain has been shown to abrogate neuroinvasiveness suggesting the potential role of translation from this region. Together, our study sheds light on distinct mechanisms that modulate JEV translation with likely consequences for viral pathogenesis.

## Introduction

Endemic to Asia-Pacific region, Japanese encephalitis virus (JEV) is the leading cause of viral encephalitis in humans and swine. Despite the availability of a vaccine, it remains a major challenge for public health and agricultural economy [1]. With ~ 11kb long positive-sense, single-stranded RNA (+ssRNA) genome flanked by untranslated regions (UTRs), JEV encodes a single multi-pass transmembrane polyprotein of about 3400 amino acids (a.a) which is proteolytically cleaved into individual structural (capsid, membrane, envelope) and non-structural (NS1, 2A/B, 3, 4A/B, 5) proteins [2]. However, translation of viral polyproteins is a complex process defined by the constraints of stoichiometry and dynamic changes in viral processes involving these proteins. Further, cotranslational folding and cleavage during polyprotein synthesis might involve regulation of ribosomal strategies like pausing and frameshifting on viral RNA (vRNA). For example, translation of JEV and closely related West Nile virus (WNV) involves ribosomal frameshifting - a process where the translating ribosome switches the protein reading frame, leading to non-uniform levels of structural and non-structural proteins [3,4]. Comparison of the vaccine strain (SA14-14-2) with the wild type (SA14) has revealed acquired mutations that are linked to attenuation of the virus [5,6]. For instance, a single nucleotide mutation (G3599A) in the vaccine strain was shown to abrogate –1 programmed ribosomal frameshifting (PRF) leading to a loss in viral neuroinvasiveness [3,7]. However, mechanistic understanding of translation strategies behind the attenuating determinants have not been entirely elucidated.

Deep sequencing ribosome-bound mRNA footprints provides a snapshot of the global translatome with the average ribosomal density per mRNA scaling linearly with its protein production rate [8]. The method, popularly known as ribosome profiling, has provided promising insights into translational regulation by RNA viruses ranging from temporal regulation of host gene expression upon Dengue virus (DENV) [9] and Hepatitis C virus (HCV) [10] infection to previously uncharacterised alternate ORFs in Coronaviruses like Mouse Hepatitis virus (MHV) [11] and SARS-CoV-2 [12]. Few RNA viruses like MHV [11], Zika virus (ZIKV) [13] and Echovirus 7 [14] also display upstream ORFs with physiological implications of the encoded peptide in virus release [14]. Additionally, the method’s accuracy at single-nucleotide resolution identifies the translation frame and has been applied to estimate actual *in cellulo* ribosomal frameshifting efficiencies during RNA-virus infections [11,12,15].

Here, we employ ribosome profiling [8] to characterise mechanisms that modulate the translation efficiency of JEV proteome during cellular infection. Using translation inhibitors *i.e* harringtonine (initiation) and cycloheximide (elongation) to stall ribosomes on viral RNA, our study reveals three key features of JEV translation: 1) —1 PRF during NS1′ synthesis results in ~ 40% reduction of ribosomes translating downstream polyprotein, 2) Significant modulation in levels of a distinct subset of ribosome-bound tRNAs that cannot be explained by virtue of codon usage and 3) Translation from an upstream ORF (uORF) using a non-canonical initiation codon in the 5′ UTR region of JEV. These events signify several strategies of translational regulation during viral polyprotein synthesis along with features which could aid the virus in neuroinvasion. Overall, our findings display the potential of translation governing factors in +ssRNA viruses’ pathogenesis by evaluating their molecular underpinnings.

## Materials and Methods

### Cells and virus

Neuro2a cells were maintained in Dulbecco’s modification of Eagle’s medium (DMEM, Himedia) supplemented with 10% (v/v) foetal bovine serum (FBS, Gibco) and 1% (v/v) Penicillin-Streptomycin (Sigma). Cells were grown to 70% confluency in 10cm dishes and infected with 5 plaque forming units (PFU) per cell of JEV P20778 Vellore strain (*AF*080251) or Dimethyl sulfoxide (DMSO, mock) in plain DMEM. After 60min at 37°C, inoculum was removed and monolayer was extensively washed with phosphate buffered saline (PBS) before replacing with complete media (DMEM with 10% FBS).

### Drug treatment and monosome purification

At 18h post infection (pi), cells were treated with either cycloheximide (Sigma-Aldrich, 100*μ*g/ml, 5min) or harringtonine (LKT laboratories, 2*μ*g/ml, 2min) followed by cycloheximide treatment for 5min. Cells were rinsed with ice-cold PBS containing 100*μ*g/ml cycloheximide. Dishes were submerged in a liquid nitrogen reservoir for 10s followed by scraping over dry ice in polysome lysis buffer (20mM Tris-HCl pH 7.5, 150mM NaCl, 5mM MgCl_2_, 1mM DTT, 1% (v/v) Triton X-100, 100*μ*g/ml cycloheximide and 25U/ml TURBO DNase *I* (Life Technologies)). Cells were collected and triturated with a 26G needle 20 times and clarified by centrifugation (13000g, 20min, 4°C). Supernatant was collected and treated with 100U/*μ*l of RNase I (Ambion) for 1h at room temperature with gentle mixing followed by inactivation with 40U of SUPERase-In RNase inhibitor (Ambion). Cell extracts were passed through Sephacryl S400 spin columns (GE) after pre-equilibration with polysome lysis buffer and ribosome-bound mRNA eluates were collected by centrifugation at 600g, 1min, 4°C [16].

### Library preparation

RNA was extracted from eluates and total lysate using TRIzol reagent (ThermoScientific). For RNA-Seq, total RNA was fragmented using 10x fragmentation reagent (Ambion) according to manufacturer’s protocol. Libraries were prepared according to original protocols of Ingolia and colleagues [17] with minor modifications. Briefly, RNA was resolved over 15% polyacrylamide TBE-urea gel using electrophoresis and a broad range of fragments (25 - 70 nts) were purified from the gel to capture both ribosome bound mRNA and tRNA segments. RNA fragments were further dephosphorylated using T4 polynucleotide kinase (NEB) for 1h at 37°C followed by heat inactivation for 10min at 75°C. Fragments were ligated to microRNA preadenylated linker (NEB) using T4 Rnl2(tr) ligase (NEB) for 2.5h at room temperature. Ligated products were size selected and purified from PAGE gel followed by reverse transcription [17] using SuperScript *III* (ThermoScientific) and eliminating RNA by NaOH hydrolysis for 20min at 98°C. cDNA is again size selected on a denaturing PAGE, gel purified and circularised using CircLigase (Epicentre) for 1h at 60° C followed by heat inactivation for 10min at 80°C. Circularised product is subjected to two rounds of rRNA depletion using biotinylated oligos [17] and streptavidin-coated magnetic beads (NEB) according to manufacturer’s protocol. The rRNA-subtracted circular product is subjected to a final round of PCR amplification with Illumina adaptor primers [17] using Phusion polymerase (NEB). All the libraries were then quantified, and quality checked by Genotypic technology services using qubit fluorimeter, real time qRT-PCR and bioanalyzer followed by sequencing on Illumina NextSeq 1 x 75 SE platform with ~ 50 and ~ 5 million reads per Ribo-Seq and RNASeq sample, respectively. Sequencing data have been deposited in ArrayExpress (http://www.ebi.ac.uk/arrayexpress) under the accession number E-MTAB-9352.

### Computational analysis

Raw sequences were quality filtered and adaptors were trimmed using FASTX-Toolkit [18]. Reads ≥ 24 nt were mapped to *Mus musculus* rRNA (Accession numbers NR003278, NR030686, NR003279, NR003280, NR046233, GU372691) and tRNA (gtRNA database) using Bowtie2 (very-sensitive-local alignment) [19] with a maximum of one mismatch and sorted into separate files. Unmapped reads were aligned to JEV strain P20778 (*AF*080251) and *Mus musculus* Refseq RNA database (mm10) without any mismatches for analyzing ribosomal footprint distribution. P-site offsets were determined by metagene analysis of host mRNA using ribogalaxy tool [20] for corresponding footprint lengths. Reads aligning to JEV were further mapped to single nucleotide by setting respective P-site offset (+12 for 28 – 30nts and +13 for 31nt). Individual fragment lengths were checked for triplet periodicity by autocorrelation of ribodensities. Footprint lengths of 28 — 29 nts for HAR-CHX and 29 – 31 nts for CHX were chosen to build subcodon riboprofiles of JEV RNA. For consistency, RNASeq profile was also constructed using +12nt offset from 5′ end of the reads. Both Ribo- and RNA-Seq counts were normalised to sequencing depth based on total number of viral mapped reads and expressed as reads per million mapped reads (RPM). Frameshift efficiency was calculated by comparing RPF densities (RPFs/gene length) before and after the frameshift region *i.e*. 180nt downstream of polyprotein start codon to 180nt upstream of frameshift start site in NS1 (276 to 3374 nts) and 180nt downstream of frameshift stop site to 180nt upstream of polyprotein stop codon (3871 to 10214 nts), respectively. Due to association of vRNA with replication and packaging during late infection stages, frameshift efficiency was not accounted for RNA levels [15]. tRNA mapping was executed using sports1.0 [21] with no mismatches and default parameters for cytoplasmic and mitochondrial tRNAs. As mitochondria employ an independent translation system and serves as internal control for relative quantification [22], individual cytoplasmic tRNAs were first normalised to total mitochondrial tRNA levels and further quantile normalised to derive relative fold changes between the samples. Sequence comparison of 5′ UTR from various JEV strains was carried out using kalign with default parameters [23]. Statistical and correlation analyses were performed using in-house written scripts.

### Construction of reporter constructs & gene expression analysis

A dual reporter vector, pCMV-sLuc-IRES-GFP [24], was employed to validate expression activity from upstream and polyprotein ORFs using cap-dependent and promoter driven expression of secretory *Gaussia* luciferase (GLuc) along with an in-frame IRES-dependent expression of green fluorescent protein (GFP) for normalisation. PCR amplicons spanning 5′UTR along with first two codons (1 - 101 nts) were generated using following primers-uORFrev (TTGAATTCGTCATGGTTATCTTCCGTTCT), ppORFrev (TTGAATTCAGT-CATGGTTATCTTCCGTTCT) and fwd (TTGCTAGCAGAAGTTTATCTGTGTGAACTTC) followed by directional cloning between *NheI* and *EcoRI* restriction sites of pCMV-sLuc-IRES-GFP. Sequence confirmed constructs were transiently transfected with 500ng of plasmid DNA using Lipofectamine 3000 (ThermoScientific, manufacturer’s protocol) in uninfected and JEV infected (24h) N2a cells seeded in 24-well plates at 0.5 * 10^6^ density per well. 24h post transfection, cell supernatants were used to monitor GLuc levels in Nunc 96-well optical-bottom plates (ThermoScientific) using Dual-luciferase reporter assay system (Promega) as per manufacturer’s protocol while the harvested cell lysates were assayed for GFP levels in Varioskan microplate reader (Thermo Fisher Scientific).

## Results

### Analysis of JEV ribosome profile

To assess the range and efficiency of JEV translation, we infected Neuro2a cells with JEV and generated translatome (Ribo-seq) and transcriptome (RNA-seq) libraries at 18 hpi. Deep sequencing and mapping of the resultant reads to viral and mouse genomes showed that vRNA represented ~ 25% of translating mRNA pool (data not shown). A comparison of nuclease protected fragment length distributions indicates ~ 40% of transcriptome aligned reads corresponding to eukaryotic ribosomal footprint size of 28 – 30 bases [8](Fig.1A). The observed minor differences in fragment length enrichment across the different Ribo-seq libraries could be attributed to technical biases during nuclease treatment. To further validate the origin of protected RNA sequences, we estimated the autocorrelation in read density for different nuclease protected fragment lengths. This revealed 28 – 31 nucleotide (nt) long reads exhibiting strong positive autocorrelation with 3-nucleotide phasing on the canonical frame (Frame 0) of the positive strand (+) JEV RNA (Fig.1B). Hence, using the P-site positions of 28 – 31 nt long ribosome protected fragments (RPFs), we built a sub-codon resolution riboprofile of JEV RNA.

**Figure 1.**
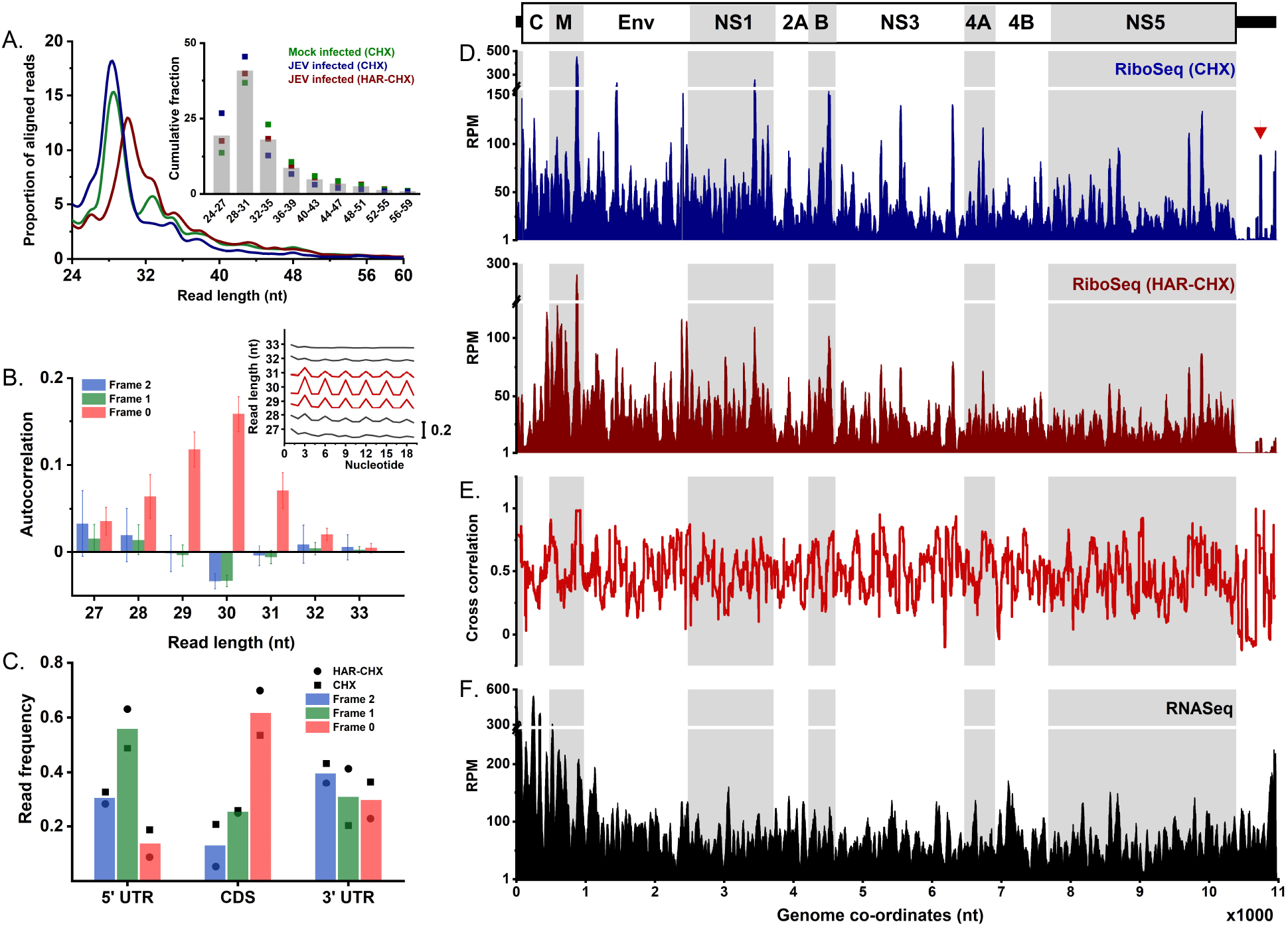
Riboprofile of JEV RNA. **A**, Read length distribution of nuclease-protected fragments compared between mock-infected and 18h JEV infected N2a cell extracts. Inset demonstrates cumulative profile of the same for every 4nt interval. **B**, Autocorrelation of ribodensities in the three frames compared between nuclease-protected fragment lengths shows 28-31 base long reads exhibiting triplet periodicity on (+)JEV RNA. Raw autocorrelation values near JEV start codon are shown in the inset. **C**, Densities of vRNA RPF P-site positions in respective frames normalised to gene length. Histograms of Ribo-Seq (**D**) and RNASeq (**F**) reads per million mapped reads (RPM) marked with P-site positions and smoothed with a 30nt sliding window across the viral genome. Arrow corresponds to nuclease protection near position 10759nt in 5′ DB region of 3’ UTR. **E**, Pearson values of cross-correlation between Ribo-Seq profiles of CHX and HAR-CHX datasets with 60nt running mean filter.

Positional arrangement of RPFs between cycloheximide (CHX) and harringtonine-cycloheximide (HAR-CHX) treated cells displayed significant and comparable nuclease protection profiles with an average cross-correlation of 0.47 over the entire coding sequence (CDS, Figs.1D and 1E). 55 – 60% of the RPFs mapped to the polyprotein reading frame (0) of JEV CDS (Fig.1C). The 5′ UTR, on other hand, harbors few nuclease protected footprints as well as shows 55% enrichment in frame 1 probably due to uORF expression (discussed later). But reads in 3’ UTR demonstrate a strong depletion immediately after the stop codon further suggesting capture of ribosomal footprints in our datasets. Residual reads at the 3’ terminus also fail to exhibit triplet phasing suggesting spurious nuclease protection due to higher order RNA structures or protection by bound ribonucleoproteins (RNPs) as the likely cause of such reads (Fig.1C). For instance, both CHX and HAR-CHX riboprofiles display a sharp increase near repeated conserved sequence 2 (RCS2) region of 5′ dumbbell-like (DB) secondary structure (position 10759 nt, marked with red arrow in Fig.1D). Sequencing of the total RNA extracted from the cell lysates revealed ~ 3-fold higher (+)RNA levels across the first 1000 nucleotides encompassing capsid and membrane encoding regions (Figs.1F and 2B). However, contrary to earlier reports [25,26], we find no evidence of subgenomic flaviviral RNA accumulation near 3’ UTR (Fig.1F).

**Figure 2.**
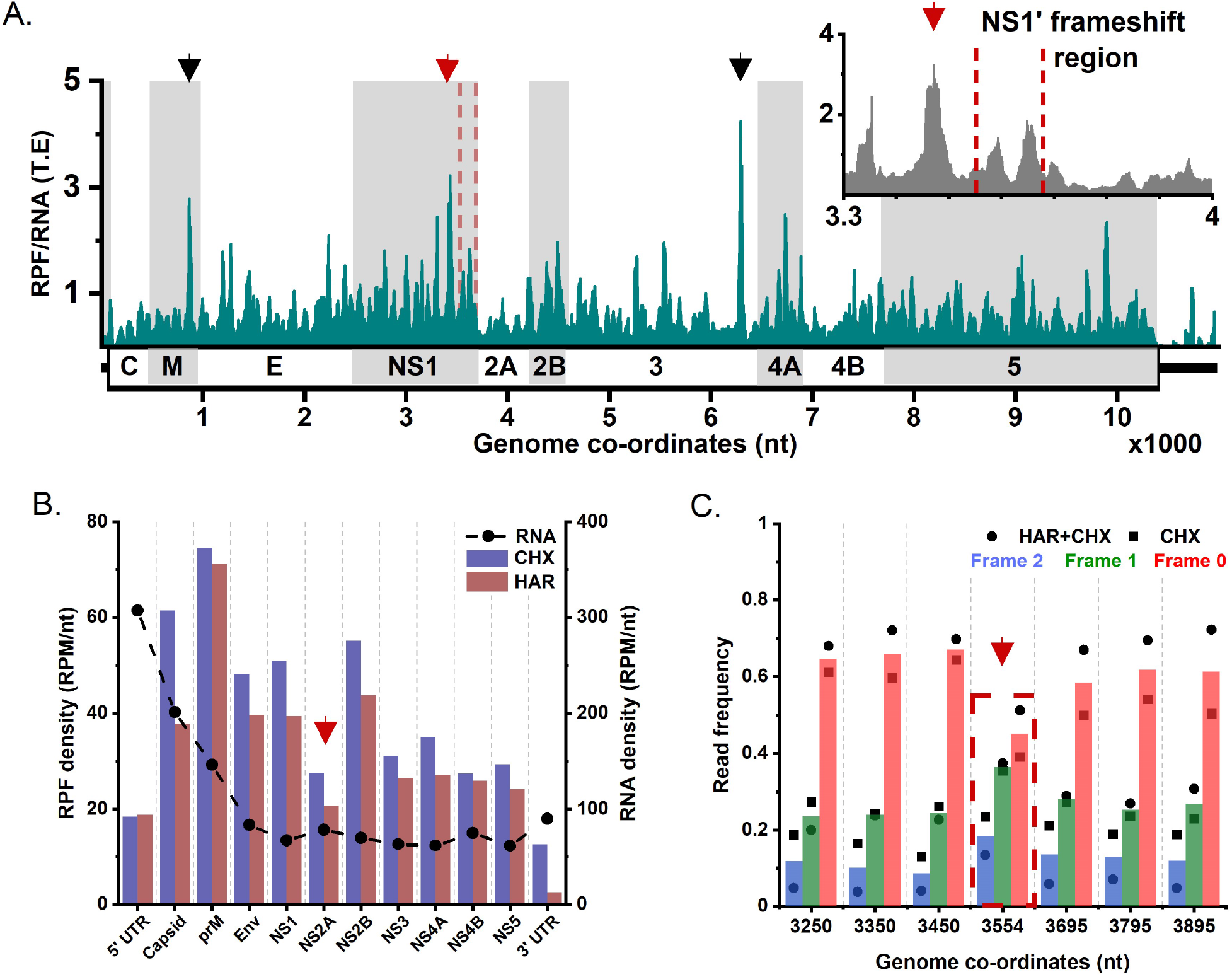
Analysis of ribosomal pausing and frameshifting on JEV RNA. **A**, Average translation efficiency (T.E) from CHX and HAR-CHX datasets compared across JEV genome with 30nt running mean demonstrates positional arrangement of ribopause sites (marked with arrows). Inset displays enlarged view of translation efficiency in NS1-2A region indicating no significant ribopauses in frameshift region (marked with dashed lines) but significant pausing 100nt upstream to frameshift site (red arrow). **B**, RPF (columns) and RNA (scatter) densities of individual viral proteins with the drop in RPF density by 40% after frameshift region shown using red arrow. **C**, Phasing of ribodensities in 3 reading frames compared near –1 PRF region (marked with red arrow).

### Impact of ribosomal frameshifting and pausing on viral protein translation

Next, we evaluated the impact of ribosomal frameshifting (–1 PRF) on relative translation efficiency (T.E) of viral proteins by normalising Ribo-seq with RNA-seq reads. A comparison across the coding region suggests comparable translational efficiencies interspersed by ribosomal pauses (spikes, Fig.2A). Such ribopauses are expected for encoding transmembrane polyproteins that undergo co-translational cleavage by host and viral proteases [27,28]. Although the overall viral translation efficiency ranks with moderately upregulated host gene expression at 18hpi (mean+S.D., data not shown), precise estimates cannot be derived as the viral genome is also engaged with other lifecycle processes like replication and packaging, which will result in an under-estimation of the translation efficiency. We therefore compare RPF densities normalised to CDS length of individual viral proteins and observe a sharp decline (by almost 40%) after the frameshift site at NS1 C-terminus, with the exception of NS2B possibly due to low ribosomal velocities (Fig.2B). This frameshift-associated drop in RPF density is supported by phasing of elongating ribosomes in the vicnity of NS1 C-terminus. Drop in frame 0 reads with a corresponding increase in frame 1 reads while encoding NS1′ frameshift region compared to frame-wise read densities immediately upstream and downstream is consistent with –1 PRF near the NS1 terminus (Fig.2C). With similar estimates reported from WNV (30 - 50%), this frameshifting can result in higher levels of structural proteins and will lead to deviations suggested in viral polyprotein stoichiometry [4]. However, considering the involvement of vRNA in parallel lifecycle processes, accurate estimates of T.E for individual viral proteins remains to be evaluated. Since a conserved RNA pseudoknot structure is shown to stimulate –1 PRF on a slippery heptanucleotide motif and generate NS1′ in JEV [3], we also scan for frameshift-associated pausing near the frameshit site. Although no ribopausing was observed at the slippery heptanucleotide sequence, we detect significant accumulation of RPFs ~ 100 nt upstream to the frameshift site (3446 nt, Fig.2A inset). This could either be a consequence of higher representation of charged and proline residues near this region or limited nuclease accessibility due to closely stacked ribosomes upstream to frameshift site. In addition, we also observe significant pause sites in NS3 (6318 nt, Asn) and membrane CDS (892 nt, Ala). However, these sites do not represent commonly associated ribosomal pause codons (eg. proline and glutamate [29]), suggesting alternate mechanisms contributing to translational pausing on vRNA.

### tRNA dysregulation revealed by ribosome-associated tRNA sequencing

Studies on RNA viruses have suggested adaptations in codon usage of viral genes to the host translation [30]. This coupling is modulated by tRNA concentrations of specific codons and regulates ribosome elongation dynamics as well as co-translational folding of proteins [28,31]. In order to understand the differential representation of stalling codons on vRNA, we first compared the global levels of ribosome-bound cytoplasmic and mitochondrial mRNA and tRNA in N2a cells upon infection. Surprisingly, cytoplasmic tRNAs exhibit ~ 6-fold higher levels upon JEV infection while all other ribosome associated RNA levels remain unperturbed (Fig.3A). As tRNAs contain modified bases which generate truncated cDNAs during reverse transcription step of library preparation [32, 33], we perform stringent alignments to quantify ribosome-protected tRNAs [21] (see methods). The tRNA levels agree well between CHX and HAR-CHX treated infected lysates (*R*^2^ = 0.97, Fig.3B) suggesting that our results are independent of procedure or protocols used for ribosome associated tRNAs [25, 33]. A subset of tRNAs also represented at higher abundance in ribosome associated fraction upon JEV infection (Figs.3B and 3C). These include glutamic acid (UUC), serine (CGA, GCU, UGA), histidine (GUG) and leucine (CAG) tRNAs (Fig.3C). Such high levels of 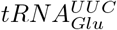 could signify a strong enrichment of GAA codon which is reported to be associated with ribopause sites in mammalian translatomes [29]. In addition, these enriched ribosome-bound tRNAs display a Pearson correlation of 0.6 against codon usage of JEV and highly expressed host genes at 18hpi (data not shown). Such correlation levels are expected from a translating transcript exhibiting ribopausing events and are consistent with ribosome-embedded yeast tRNA profiles generated using targeted tRNA library preparations [25,33]. Since mice infected with JEV and WNV show upregulation of certain aminoacyl tRNA synthetases [34], it is possible that translation efficiencies dynamically change as a function of local tRNA availability. This has important implications for the regulation of viral translation and thereby, viral growth dynamics in the cell.

**Figure 3.**
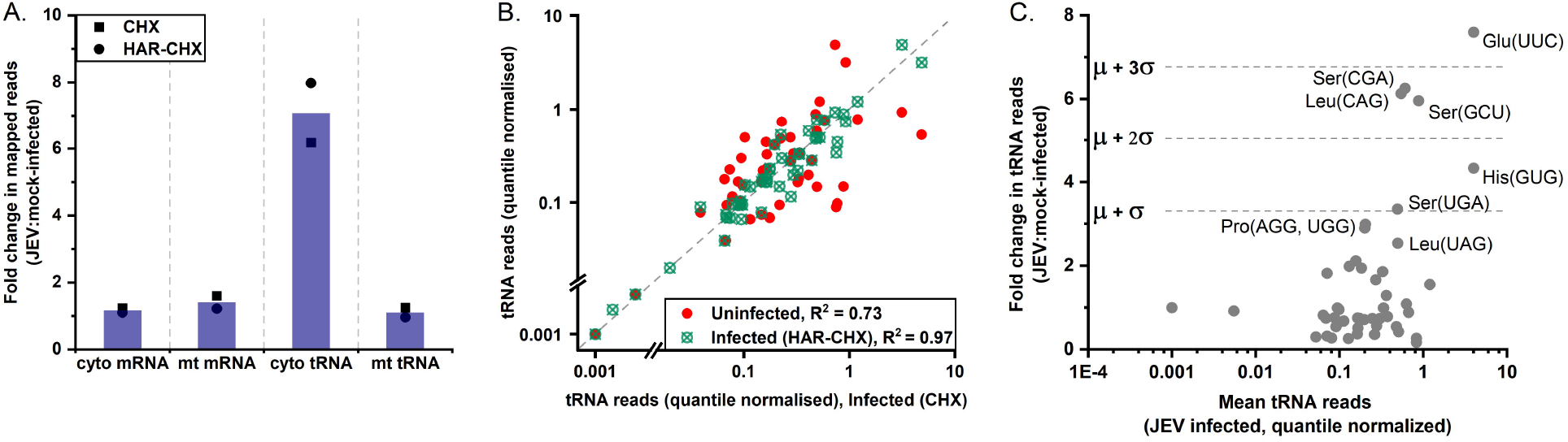
JEV infection causes selective recruitment of tRNAs to ribosomes. **A**, Fold change in total aligned reads of ribosome-bound mRNA and tRNAs from cytosol and mitochondria between mock-infected and 18h JEV infected N2a cell extracts. **B**, Scatter plot between normalised levels of ribosome-bound cytosolic tRNAs in CHX-infected sample against HAR-CHX infected (R^2^ = 0.97, green) and mock infected lysates (R^2^ = 0.73, red). **C**, Relative fold changes in cytosolic, ribosome-bound tRNA normalised to total mitochondrial tRNA reads. Significant threshold levels in fold changes at 68% (mean+S.D, *μ* + *σ*), 95% (*μ* + 2*σ*) and 99% (*μ* + 3*σ*) are represented using dashed lines.

### Translation of upstream ORF in 5′ UTR

We next examine the observed nuclease protection profile and phasing of ribosomes in 5′ UTR (Figs.1C and 4A). Both CHX and HAR-CHX treated riboprofiles show distinct reads from 38^*th*^ nt which harbours a non-canonical start codon-UUG in frame 1 (Fig.4A). The putative uORF would generate a 22 a.a long peptide (LRRIERLVQFKQFFRTEDNHD) with a termination codon immediately downstream of canonical start codon (Fig.4C). A recent study on riboprofile of ZIKV also showed ribosomal initiation scanning from out-of-frame non-canonical start codons present in the 5′ UTR - CUG (uORF1) and UUG (uORF2) [13]. However, unlike ZIKV riboprofile where initiating ribosomes tend to stall more at uORF start codons compared to AUG, riboprofile of JEV (CHX) in N2a cells shows that the RPF density is ~ 3.2x higher at the canonical start codon compared to the uORF start site (Figs.4A and 2B). Translation initiation from both JEV start codons was confirmed by luciferase-based translation reporter constructs (Fig.4B). Consistent with our ribosome profile findings, polyprotein ORF (ppORF) expresses 2.7 – *4x* more efficiently than uORF (Fig.4C), as expected from poor initiation context of UUG [29]. Interestingly, JEV infection appears to stimulate expression from UUG start site by almost 67% suggesting viral or virus-induced host *trans*-regulatory factors promoting uORF translation (Fig.4D). Upon sequence comparison of 5′ UTR across JEV strains, the alternate start codon exhibits 100% conservation except for the attenuated strain-JEV SA14-14-2 harbouring U39A mutation and generating a previously overlooked stop codon (Fig.4E). Implications of this disrupted alternate start site in the vaccine strain remains to be evaluated.

**Figure 4.**
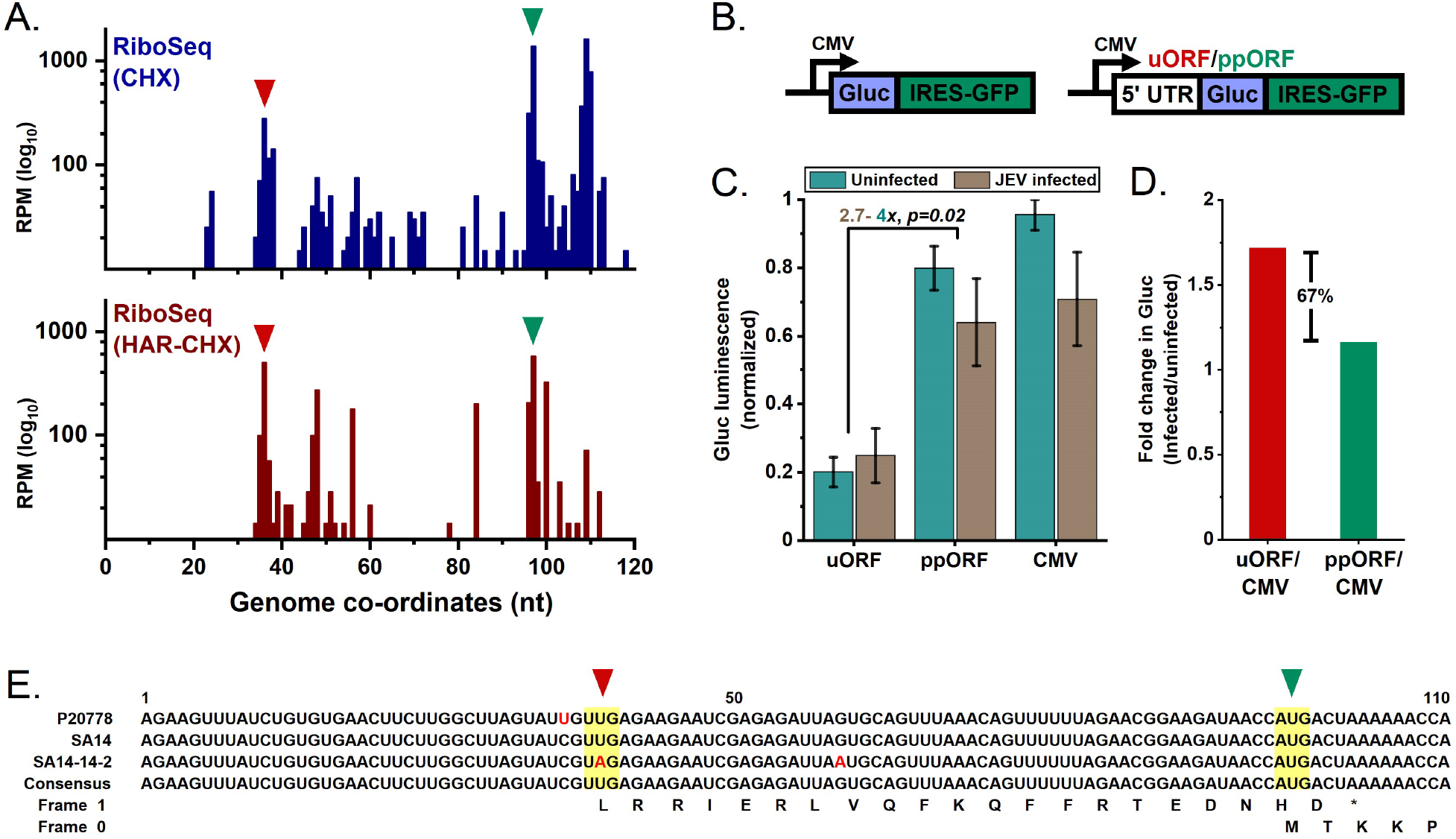
uORF translation in JEV. **A**, Riboprofile of JEV 5′ UTR from CHX and HAR-CHX treated samples reveals ribosomal initiation scanning from an alternative start codon-UUG at 38^*th*^ nt in frame 1. Arrows denote position of start codons. **B**, Schematic of reporter constructs (pCMV-sLuc-IRES-GFP, left)) used to validate uORF and ppORF expression from JEV 5′ UTR (right). **C**, Normalised *Gaussia* luciferase (Gluc) levels demonstrating expression from UUG (frame 1) and AUG (frame 0) of JEV 5′ UTR. Data is represented as mean±s.e.m. for n=4 biological replicates and p-value estimated using Student’s t-test. **D**, Fold change in Gluc expression levels from uORF and ppORF compared upon JEV infection in N2a cells relative to pCMV expression. **E**, Sequence comparison of JEV 5′ UTR wild type consensus with the vaccine strain, SA14-14-2, demonstrates U39A mutation. Peptide sequences of wild type strain from respective frames are shown below nucleotide alignment and the translation start sites are highlighted and shown with arrows.

## Discussion

In this study, we utilise ribosome profiling and tRNA sequencing to investigate possible translation regulation strategies employed during JEV infection. A comparative analysis of riboprofiles with translation initiation and elongation inhibitors allowed us to examine *bona fide* RPFs exhibiting 3-nucleotide phasing. The sensitivity of this method is reflected in detection of previously characterised –1 programmed frameshift in NS1 CDS [3]. Few members of flaviviruses, like JEV and WNV, exhibit such frameshifting and express proteins from overlapping frames [4]. While our data does not show any significant pausing at the frameshift site as observed in encephalomyocarditis virus [15], the rise in ribodensity ~ 100nt upstream suggests likely modulation of ribosome progressivity. Nonetheless, the reduction in actively translating ribosomes leads to almost 40% drop in downstream non-structural proteins (NS2A–NS5). These proteins play an important role in establishing replication milieu in host cell via membrane re-organization. Several RNA viruses sequester polymerase (NS5) away from the site of synthesis *i.e* membranous vesicles, and also regulate viral protein stoichiometry spatially post translation [35, 36]. Therefore, modulation of NS2A - NS5 levels during translation via ribosomal frameshifting presents a critical point in infectivity for regulating the extent of membrane re-organization and vRNA replication.

Our analysis also sheds light on how ribosomal pausing and tRNA regulation can impact translational efficiencies and possibly modulate polyprotein folding and processing. We report major pause sites across JEV genome with the most significant ones in membrane, NS1 and NS3 CDS while the representative pause codons are distinct from those reported in mammalian systems [29]. We also identify a subset of ribosome associated tRNAs whose levels are modulated globally upon JEV infection (Fig.3). However, such tRNA abundances remain to be critically examined as recent studies indicate interference of cycloheximide in quantifying bound tRNA fractions due to ribosomal conformational locks [31]. Nevertheless, this discriminating tRNA subset during JEV infection could provide a possible antiviral intervention strategy. For example, a recent study demonstrated impairment in WNV infectivity upon depletion of schlafen-11 which prevents WNV-induced changes in a tRNA subset translating 11.8% of viral polyprotein [37]. Also, schlafen-11 was shown to bind tRNAs essential for HIV protein synthesis during later stages of infection in a codon usage dependent manner [38]. It is additionally possible that these tRNA might be involved in translation-independent processes like viral replication (eg. retro- and bromoviruses [39]) or novel functions with their site-specific interactions on viral genome (eg. 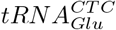 and 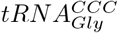 with ZIKV [40]). Although studies have indicated adaptation in viral codon usage to host organism or tissue [41,42], our findings suggest its re-evaluation in the context of the anticodon counterpart (tRNA) dynamics that will arise due to virus-induced cellular rearrangements. Manipulating virus codon usage to attenuate the virus has recently shown promising advancements in vaccination of mice models [43–47]. Combined with tRNA perturbation, efficacy of these putative attenuated vaccine targets can be further improved.

Despite the employment of translation inhibitors to enrich ribosome associated fragments, both UTRs display unique nuclease protected regions. While reads in 3’ UTR fail to exhibit phasing, RCS2 region of 5′ DB displays significant nuclease protection. This region, arising from duplication of CS2 region in 3’ DB, is observed across JEV and DENV serotypes and mutations in CS2 region have also shown to decrease translation and replication in the latter [48]. Despite the absence of a polyA tail, it was shown that A-rich sequences flanking these DB structures interact with polyA binding protein (PABP) and enhance translation by serving as a circularisation factor [49]. We speculate that CS2 and RCS2 duplicate sequences in 3’ UTR facilitate a similar interaction and possibly modulate vRNA translation by circularisation and ribosome recycling. We also observe a striking nuclease protection profile in 5′ UTR upon comparing CHX and HAR-CHX datasets which exhibit characteristics of an uORF (Fig.4A). Translational activity of the uORF was validated using reporter constructs (Fig.4B). While the functional significance of the putative peptide remains unknown, a close overlap of uORF stop codon (100^*th*^ nt) with main ORF’s start codon (96^*th*^ nt, Fig.4C) could directly impact ribosome loading and usage rates-key parameters in regulating protein synthesis [50]. Although a canonical structure of 5′ UTR is shared amongst flaviviruses [51], this strategy of translation initiation might suggest the enhanced translational efficiency of JEV and ZIKV [13] over DENV [9]. This site exhibits high conservation across JEV strains with the exception of vaccine strain (SA14-14-2) harbouring a silent mutation. A reverse genetic approach incorporating U39A mutation in wild type JEV SA14 backbone reduced mice neuroinvasiveness but not neurovirulence of the virus with no significant differences in viral titres under *in vitro* conditions [5]. However, it remains to be tested whether this intervention is a consequence of the putative short peptide encoded from the uORF or a *cis* acting element important for RNA transport as suggested in tick borne encephalitis virus [52]. A similar strategy of uORF and polyprotein expression is employed by enteroviruses where the uORF encoded protein was shown to facilitate virus growth in gut epithelial cells [14]. This remarkable display of tropism using short uORFs, along with other *cis* acting elements, by RNA viruses further adds to the purifying selection imposed on UTRs. In summary, our findings on translation strategies highlight new insights that can serve as perturbation nodes during JEV infection cycle, paving new avenues for therapeutic interventions against neurotropic flaviviruses.

## Acknowledgments

The authors thank Prof. P.N. Rangarajan (BC, IISc) for providing JEV P20778 virus and Neuro2a cells; Prof. Udaykumar Ranga (MBGU, JNCASR) for the provision of pCMV-sLuc-IRES-GFP construct. We are also indebted to Dr. Sumanta Mukherjee (IBM, Bengaluru) for discussions and helping with analysis.

## Author Contributions

VAR and RR designed the experiments and wrote the manuscript. VAR performed the experiments. VAR and MS analysed the data. VAR, MS and RR interpreted the results.

